# Structural features within the NORAD long noncoding RNA underlie efficient repression of Pumilio activity

**DOI:** 10.1101/2021.11.19.469243

**Authors:** Omer Ziv, Svetlana Farberov, Jian You Lau, Eric Miska, Grzegorz Kudla, Igor Ulitsky

## Abstract

It is increasingly appreciated that long non-coding RNAs (lncRNAs) carry out important functions in mammalian cells, but how these are encoded in their sequences and manifested in their structures remains largely unknown. Some lncRNAs bind to and modulate the availability of RNA binding proteins, but the structural principles that underlie this mode of regulation are underexplored. Here, we focused on the NORAD lncRNA, which binds Pumilio proteins and modulates their ability to repress hundreds of mRNA targets. We probed the RNA structure and long-range RNA-RNA interactions formed by NORAD inside cells, under different stressful conditions. We discovered that NORAD structure is highly modular, and consists of well-defined domains that contribute independently to NORAD function. We discovered that NORAD structure spatially clusters the Pumilio binding sites along NORAD in a manner that contributes to the de-repression of Pumilio target proteins. Following arsenite stress, the majority of NORAD structure undergoes relaxation and forms inter-molecular interactions with RNAs that are targeted to stress granules. NORAD sequence thus dictates elaborated structural domain organization that facilitates its function on multiple levels, and which helps explain the extensive evolutionary sequence conservation of NORAD regions that are not predicted to directly bind Pumilio proteins.

## Introduction

Mammalian genomes are pervasively transcribed, with tens of thousands of unique loci producing long RNA molecules that do not serve as templates for production of functional proteins. These RNAs, which are collectively called long noncoding RNAs (lncRNAs) (Ulitsky and Bartel, 2013), closely resemble mRNAs on the molecular level: they are capped, polyadenylated, and usually spliced. Compared to protein-coding genes (PCGs), lncRNAs as a group are somewhat shorter, are expressed at lower levels, and are more tissue-specific (Derrien et al., 2012). Whereas the catalogs of lncRNAs in mammals are well annotated, their modes of action remain largely obscure. The low abundance and large size of the lncRNA molecules make it difficult to apply the same biochemical approaches used for elucidating the molecular mechanisms of other classes of RNAs.

While most of the functionally-characterized lncRNAs act in the nucleus, many other lncRNAs accumulate in the cytosol (Derrien et al., 2012), and plausibly act in post-transcriptional regulation. Specifically, modulation of the activity of RNA binding proteins (RBPs) or small RNAs by lncRNAs via competition for binding is one of the most commonly suggested modes of action for lncRNAs to date. The molecularly indistinguishable characteristics of mRNAs and lncRNAs make the latter effective decoys in theory. However, there are substantial doubts regarding the feasibility of the direct competition model, in light of the low expression of lncRNAs compared to that of RBPs, and the consequently relatively few RBP binding sites offered by any lncRNA gene compared to the sites found throughout the transcriptome (Jens and Rajewsky, 2015). Understanding the features that turn lncRNAs into effective decoys is crucial for evaluating how common this mode of action is and for designing synthetic RNAs capable of competing off the activity of specific RBPs.

The NORAD lncRNA is one of the most abundant and conserved lncRNAs in mammalian cells (Lee et al., 2016; Munschauer et al., 2018; Tichon et al., 2016). NORAD presence is required for preventing chromosome instability in HCT 116 cells (Elguindy et al., 2019; Lee et al., 2016; Munschauer et al., 2018) as well as preventing premature aging in mice (Kopp et al., 2019). Additional consequences of loss of NORAD were described in endothelial cells (Bian et al., 2020; Zhao et al., 2020) and in cancer cells (Soghli et al., 2021). NORAD accumulates to hundreds or even thousands of copies per cell, mostly in the cytoplasm (Elguindy et al., 2019; Lee et al., 2016; Matheny et al., 2020; Tichon et al., 2016), although nuclear localization and activity were also reported (Munschauer et al., 2018). NORAD contains ∼20 Pumilio Recognition Elements (PREs), which are binding sites for PUM1/2, two members of the Pumilio family of RBPs. PUM1/2 post-transcriptionally repress gene expression, and modulation of NORAD expression levels results in a corresponding transcriptome-wide change in the abundance of Pumilio targets (Elguindy and Mendell, 2021; Kopp et al., 2019; Lee et al., 2016; Tichon et al., 2016). While the total number of sites offered by NORAD is substantially higher than that offered by any other single gene, and is comparable to the total number of PUM1/2 protein molecules in a human cell, it is still small compared to the total number of binding sites offered by all other Pumilio targets combined (Bohn et al., 2018). The formation of phase-separated NORAD-Pumilio (NP) bodies was recently shown to enable a more efficient competition for Pumilio binding by NORAD (Elguindy and Mendell, 2021).

Little is known about the functionality of the non-PRE regions in NORAD. We recently used a massively parallel RNA assay to determine sequences within NORAD, mostly found near its 5’ end, which are sufficient for effective NXF1-dependent export of an intronless RNA (Zuckerman et al., 2020). Both the PREs and non-PRE regions within NORAD were recently shown to be required for effective NORAD recruitment into stress granules upon metabolic stress (Matheny et al., 2020). Additionally, a sequence in the 5’ region of NORAD was shown to be associated with RBMX protein and to be required for genome integrity (Munschauer et al., 2018). We found that the sequence of NORAD contains 12 sequence-similar “NORAD Repeat Units” (NRUs) (Tichon et al., 2018, 2016). The Mendell lab recently showed that mutating all the PREs in NORAD is sufficient for abolishing its ability to prevent chromosome number instability in HCT116 cells, and that a sequence containing only NRUs 7 and 8 is sufficient for this activity (Elguindy et al., 2019) (NRUs 7+8 correspond to ND4 in the notations used by the Mendell lab). Still, the function of the vast majority of NORAD sequence, much of which is highly conserved among mammals, remains unknown, and it is unclear if and how it contributes to its function in antagonizing Pumilio activity.

The structure of several lncRNAs was previously interrogated *in vitro* using synthetic refolded RNA (Chillón and Pyle, 2016; Hawkes et al., 2016; Lin et al., 2018; Liu et al., 2017; Novikova et al., 2012; Somarowthu et al., 2015; Xue et al., 2016). More recently, transcriptome-wide methods provided additional structural data from within cells. These efforts resulted in important structural and functional insights for several lncRNAs (Fang et al., 2015; Lu et al., 2016; Smola et al., 2016). Yet, while *in vitro* studies cannot reflect the impact of the cellular environment on lncRNA structure, transcriptome-wide studies typically result in low coverage and low resolution per transcript. It is increasingly appreciated that inside cells, the structure of RNA is inherently dynamic (Li and Aviran, 2018; Lu et al., 2016; Morandi et al., 2021; Tomezsko et al., 2020; Zhang et al., 2021; Ziv et al., 2020, 2018). This phenomenon is best described for riboswitches, for mRNAs undergoing splicing, and for the genome of several RNA viruses, where structural plasticity increases the functional capacity of the RNA. However, whether mammalian noncoding RNAs adopt alternative conformations during their life cycle, and the functional importance of these conformations, remains under-explored.

Here we combined *in vivo* structural probing with affinity selection of a single lncRNA, resulting in high depth and high-resolution maps reflecting the folding of NORAD inside human cells. We reveal that NORAD folds into discrete structural domains, and undergoes structural reorganization in response to certain stress stimuli. NORAD structural domains cluster together the Pumilio binding sites along its sequence and facilitate the ability of NORAD to derepress Pumilio targets.

## Results

### NORAD structure in control and stressed cells

In order to probe the RNA structure of NORAD within living cells we applied the COMRADES method with probes targeting NORAD in HCT116 cells (Ziv et al., 2018). Briefly, RNA base-pairing was crosslinked inside living cells using Psoralen-TEG-Azide, after which, total RNA was extracted and NORAD was pulled down using a tiling array of antisense biotinylated probes. Following RNA fragmentation, we used click chemistry to attach biotin to the crosslinked RNA, and it was pulled down using streptavidin beads. Half of the resulting RNA - the ‘interactions’ sample - was proximity ligated, followed by a reversal of the crosslink and sequencing. In the other half - the ‘control’ sample - the crosslink was first reversed, after which the RNA was proximity-ligated and sequenced ץ

In order to characterize the structural organization of NORAD in different cellular conditions, we used untreated cells, cells treated with doxorubicin (Doxo), a DNA damaging reagent previously shown to influence NORAD abundance and subcellular localization (Elguindy et al., 2019; Lee et al., 2016; Munschauer et al., 2018), and arsenite (Ar), a reagent that leads to metabolic stress and a strong shift in the localization of NORAD to stress granules (Khong et al., 2017; Khong and Parker, 2018; Matheny et al., 2020; Moon et al., 2020). Analysis of the sequencing data showed that NORAD reads were enriched more than 1,000-fold following pulldown with biotinylated probes. Approximately 4% of all reads were chimeric and ∼1% represented the base-pairing of NORAD inside human cells. In contrast, only <0.1% of the reads in the control samples in which reverse crosslinking was performed before the proximity ligation were NORAD:NORAD chimeras, demonstrating a good signal-to-noise ratio.

We first examined the overall distribution of RNA-RNA interactions within NORAD. Replicates of untreated cells were highly reproducible, similar to those of Doxo-treated cells, and less similar to those in Ar-treated cells (**Fig. 1A**). Visual examination of the 2D interaction map revealed clusters of intramolecular interactions at the 5’ and 3’ ends of NORAD, with more focal long-distance interactions within the middle part of NORAD which harbors the Pumilio Recognition Elements (PREs). These clusters of intramolecular interactions were not present in control data obtained from cells where crosslinked reversal proceeded the proximity ligation (**Fig. S1**). Furthermore, some of the interactions coincided with the PRE regions (as discussed in detail below).

**Figure 1.**
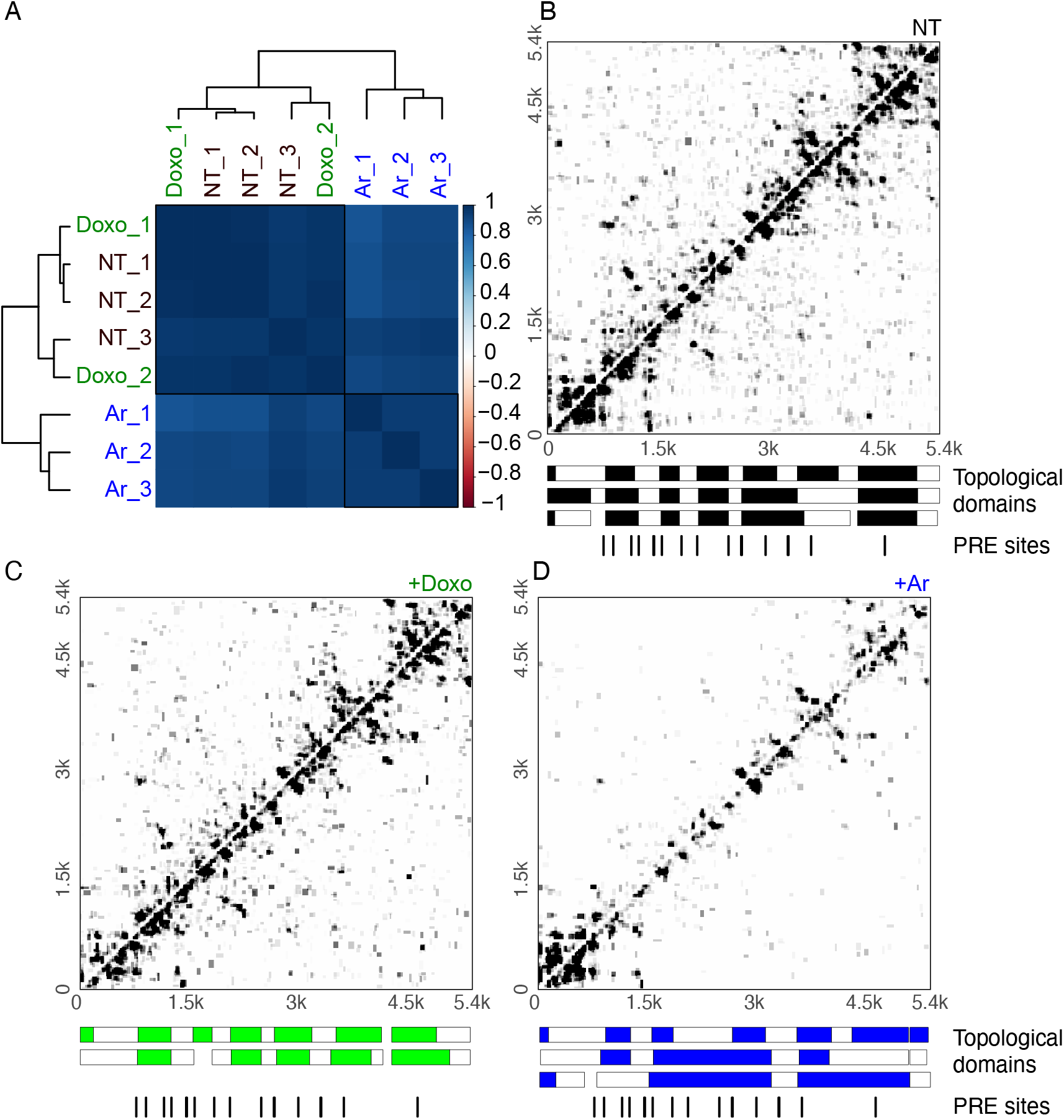
Structural organization of NORAD. **(A)** Correlation heatmap of NORAD replicates without treatment (NT_1-3), with Doxo (Dox_1-2), or Ar (Ar_1-3) treatment. Color intensity reflects the strength of correlation. **(B-D)** Domain organization of NORAD without treatment and with Doxo (C) or Ar treatment (D). Interaction maps are shown for experiments NT_1 (B), Doxo_1 (C) and Ar_1 (D). The blocks under the heatmaps show the division of NORAD into domains in each independent replicate experiment. Domains were called with the program TopDom (Shin et al., 2016) using a 300-nt window size.

We next sought to test whether NORAD folds into distinctive structural domains. We applied the TopDom algorithm, originally developed for partitioning genomes into topological domains using chromatin conformation capture datasets (Shin et al., 2016). We identified topological domains within NORAD that were highly concordant between replicates and similar between the untreated and the Doxo-treated cells (**Fig. 1C**). In contrast, Ar treatment led to a substantial reduction of contacts between most NORAD regions, with a notable exception of the 5’ region (**Fig. 1D**). Taken together, NORAD folds into distinctive spatial domains within cells and undergoes global unfolding throughout most of its sequence in response to arsenite stress.

### Dynamic changes in NORAD structure upon arsenite treatment

Analysis of chimeric reads revealed a marked reduction of intra-NORAD interactions following arsenite treatment, with a notable exception of the 5’ region, where a substantial number of interactions remained and even became more pronounced in Ar-treated cells (**Fig. 2**). In order to formally test the changes in intra-NORAD interactions upon metabolic stress, we used DESeq2 (Love et al., 2014) to compare the number of chimeras in windows of 10 nt along NORAD. This analysis showed a significant increase in contacts in the 5’ module, alongside a reduction in the middle region of NORAD, which was most pronounced within individual NRUs (**Fig. 2**, top). Inspection of the regions with the largest changes showed limited changes in the local structures (**Fig. 2**, bottom), but rather an overall reduction in interaction frequency in Ar-treated cells, suggesting the structure of the central and 3’ part of NORAD becomes globally unfolded upon Ar treatment, rather than adopting a particular alternative fold.

**Figure 2.**
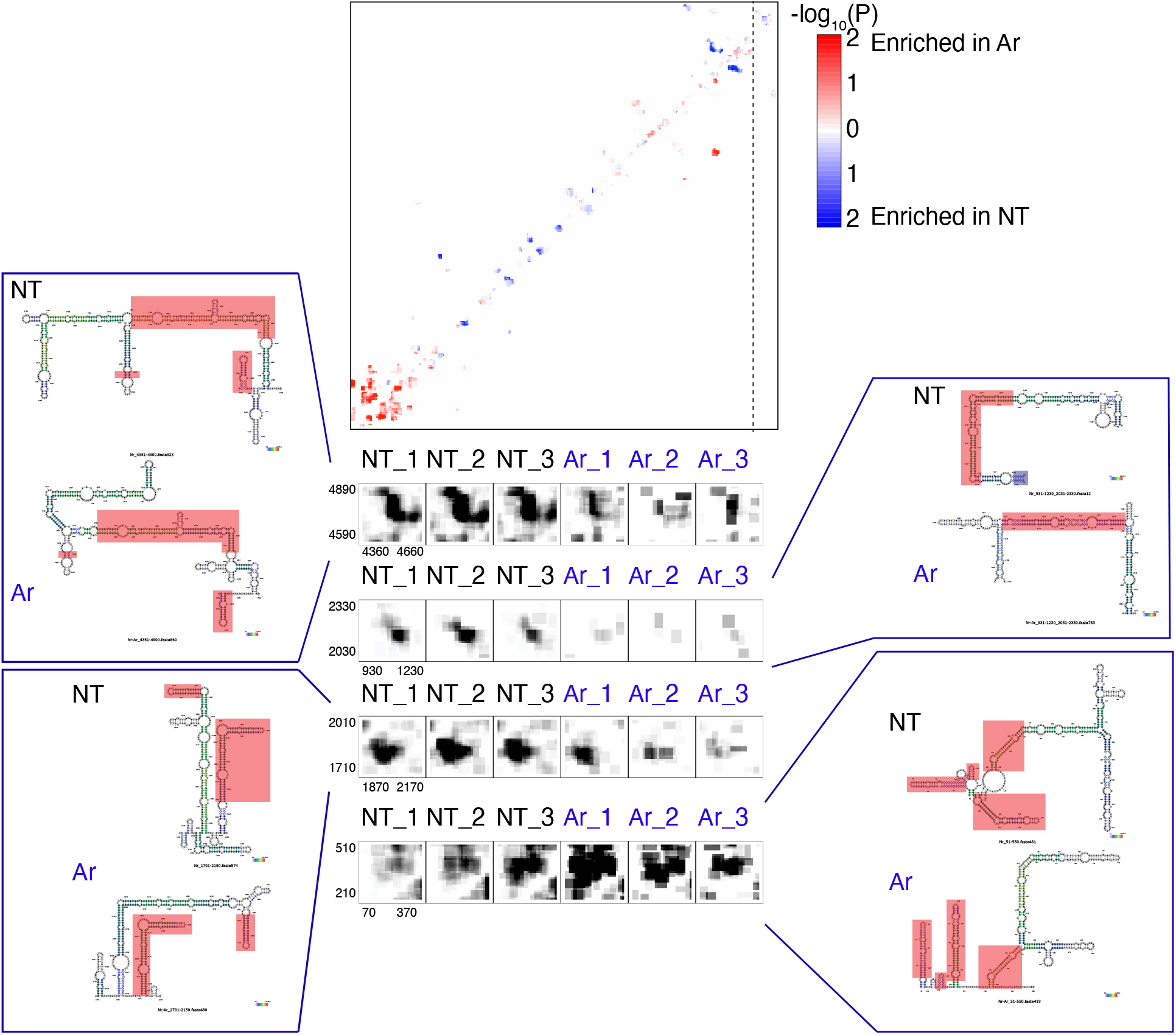
Comparison of NORAD:NORAD interactions between NT and Ar treated cells. Statistical significance of enrichment in either condition was calculated using DESeq2 in 10-nt windows. Red color indicates enrichment in Ar-treated cells; blue color, enrichment in no-treatment cells. Bottom: coverage of chimeras in selected regions in the indicated samples, with panels showing the predicted and color-coded structures based on the three NT and the three Ar replicates. Regions shaded in red have identical predicted structures in NT and in Ar-treated cells.

### Spatial clustering of NORAD PREs

We next combined all the RNA-RNA interactions together and examined the boundaries of the structural domains identified by TopDom and the numbers of inter- and intra-molecular interactions in the context of the 12 NRUs, the 5’ region preceding them, and the 3’ region (**Fig. 3**). In the following analysis, we focused on eight canonical “PRE clusters” found in alignable positions in eight of the NRUs, with each cluster containing one or two PREs. These corresponded to the most prominent peaks in the PUM1 CLIP data from HCT116 cells (Munschauer et al., 2018) (**Fig. 3**). Domains typically contained several NRUs, with two domains corresponding to the 5’ region upstream of the 12 NRUs and two containing the 3’ region downstream of them. Interestingly, regions surrounding the PREs had an overall lower tendency to form interactions with other RNA molecules (**Fig. 3**), which was evident for four of the eight PRE-containing regions in no-treatment and Doxo-treated cells and for all the PREs in arsenite-treated cells (**Fig. S2A**). Upon arsenite treatment, there was a strong reduction in the normalized number of chimeric reads for both intra-NORAD and NORAD-other interactions, which was much less pronounced in the two 5’ domains, which were overall much more G/C-rich than the rest of NORAD sequence. Interestingly, the first 1/8 of NORAD sequence, which was substantially less affected by arsenite treatment, is also the part that is least capable of recruiting a reporter RNA to stress granules upon arsenite treatment (Matheny et al., 2020).

**Figure 3.**
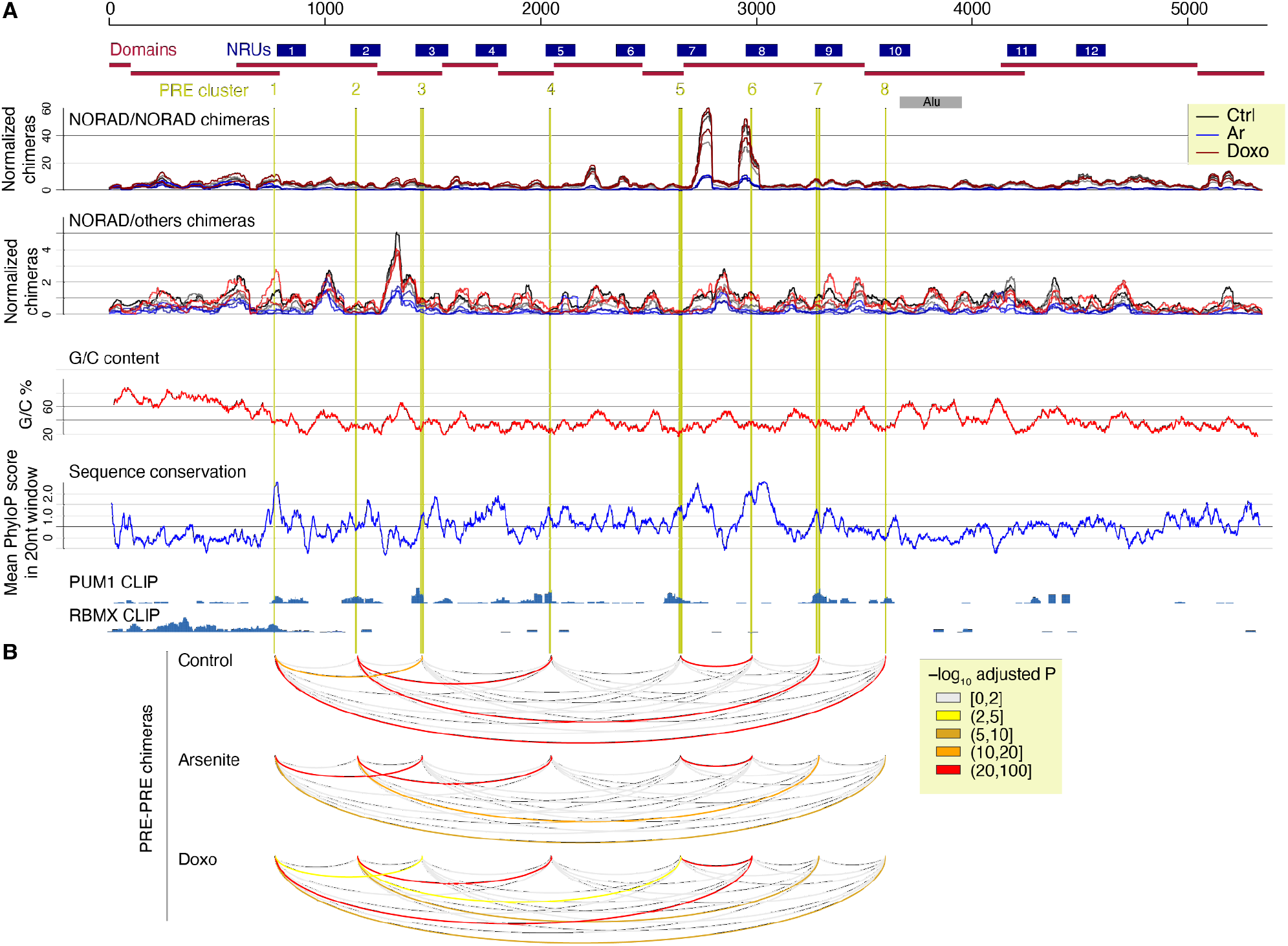
Distribution of intra- and inter-molecular RNA interactions along the NORAD sequence. (A) NORAD locus, with domains identified when considering all the chimeric reads, and NRUs from (Tichon et al., 2016) demarcated. Canonical PREs found in corresponding positions in the NRUs are marked by vertical lines. Normalized number of intra-NORAD and NORAD-other chimeric reads is shown separately for each biological replicate with the indicated color code. G/C content and PhyloP scores from the 100-way whole-genome alignment from the UCSC browser were computed for windows of 20 bases. CLIP data coverage at the bottom for PUM1 and RBMX are from (Munschauer et al., 2018). (B) NORAD arc diagram connecting pairwise combinations of PRE clusters. Colors are based on -log_10_ of Bonferroni adjusted p-values. Colored arcs signify a significantly higher number of chimeric reads in PRE-PRE cluster pairs than permuted positions in NORAD. (Arc diagram made using the “R4RNA” R package (Lai et al., 2012).

When examining the predicted structures with the strongest experimental support (**Fig. 4** and **Figs. S3-4**), we noted that five of the eight PRE clusters (1–3, 5, and 6) appeared to be in close spatial proximity to other PRE clusters, which was also evident when examining the distribution of chimeras formed by each PRE cluster, an analysis which does not rely on any explicit structure prediction (**Fig. S2B**). In order to formally test whether spatial clustering of PREs takes place, we analyzed the number of chimeric reads connecting regions ±50 nt around the PRE clusters, and compared them to random equidistant positions within NORAD. For five of the PRE pairs, the number of chimeric reads was significantly (P<0.05, adjusted for the number of PRE cluster pairs) larger than expected by chance in all experimental conditions, and two additional pairs were significant only after Doxo treatment (**Table S1)**. The overall structure of NORAD thus positions PREs in close spatial proximity to each other, in a manner that may facilitate the formation of NORAD-Pumilio bodies (Elguindy and Mendell, 2021) (see Discussion).

**Figure 4.**
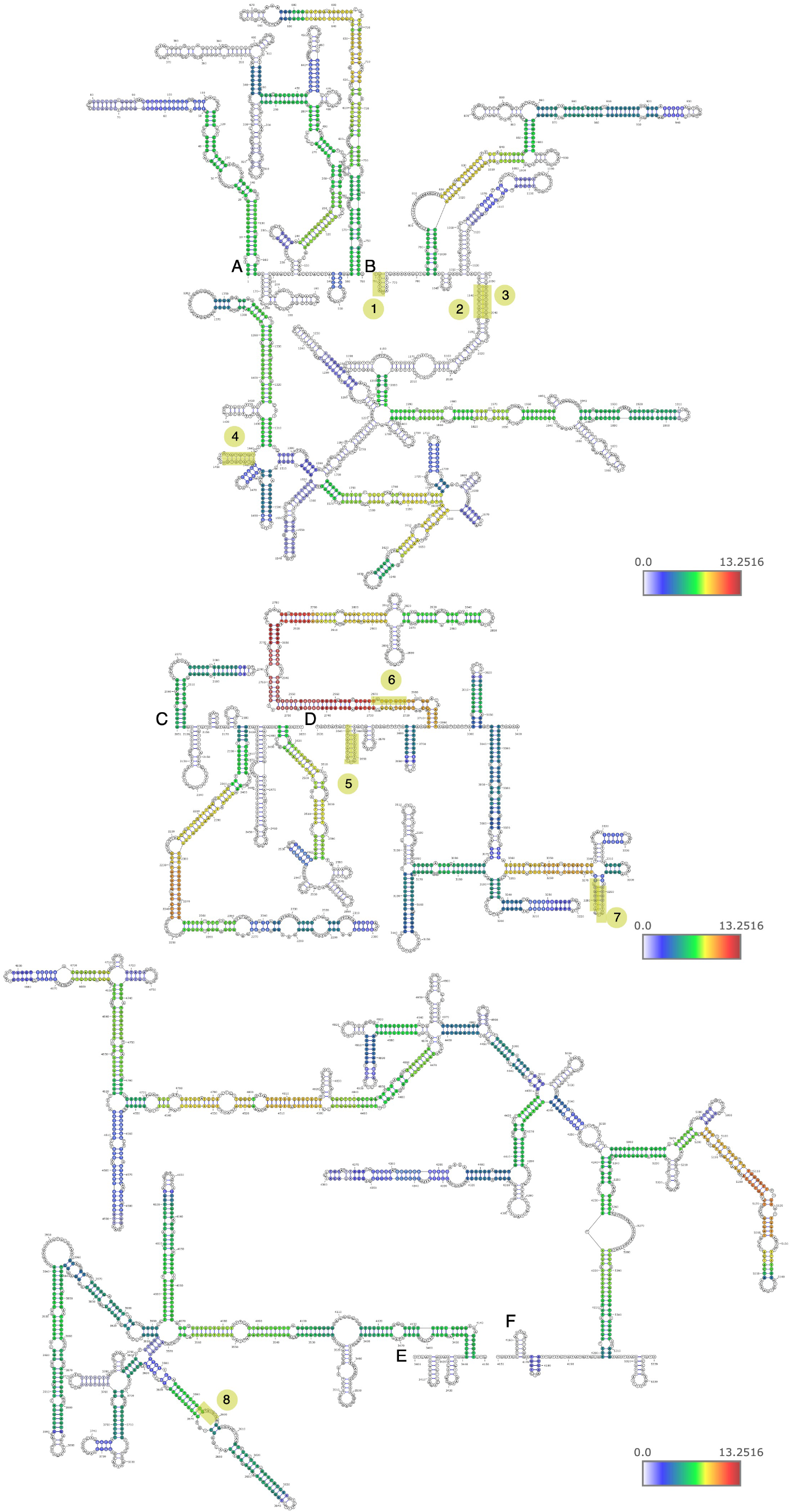
Predicted structures of NORAD domains. Predicted structures of separately folded NORAD regions 1-760 (A), 761-2050 (B), 2051-2630 (C), 2631-3400 (D), 3401-4150 (E), and 4151-5339 (F). Bases are colored based on log_2_ of the number of reads normalized to chimeric reads per million mapped reads. The eight clusters of PREs are shaded and numbered as in Figure 2.

### Contribution of the modular structure of NORAD to de-repression by Pumilio proteins

In order to study the contribution of different sequences within NORAD to its ability to inhibit Pumilio activity, we generated a reporter that contains 8 PRE elements within the 3’ UTR of Renilla luciferase (8XPRE). Firefly luciferase, used as a control, was expressed from the same vector. Expression was compared to that of a reporter where all the canonical PRE <u>UGU</u>AUAUA sites were mutated to <u>ACA</u>AUAUA, expected to abolish Pumilio binding (8XmPRE) (**Fig. S5A**). We previously used a similar reporter with 3 PRE elements and showed it was sensitive to knockdown or repression of PUM1/2 in U2OS cells (Tichon et al., 2016), and introducing the reporters into cells over-expressing different NORAD variants is an effective and quantitative way to study the efficiency of NORAD sequences in inhibiting Pumilio activity. The 8XPRE reporter was de-repressed by ∼4-fold after combined knockdown of PUM1 and PUM2, allowing for a substantial dynamic range for measurements of Pumilio repression activity (**Fig. 5A-B**).

**Figure 5.**
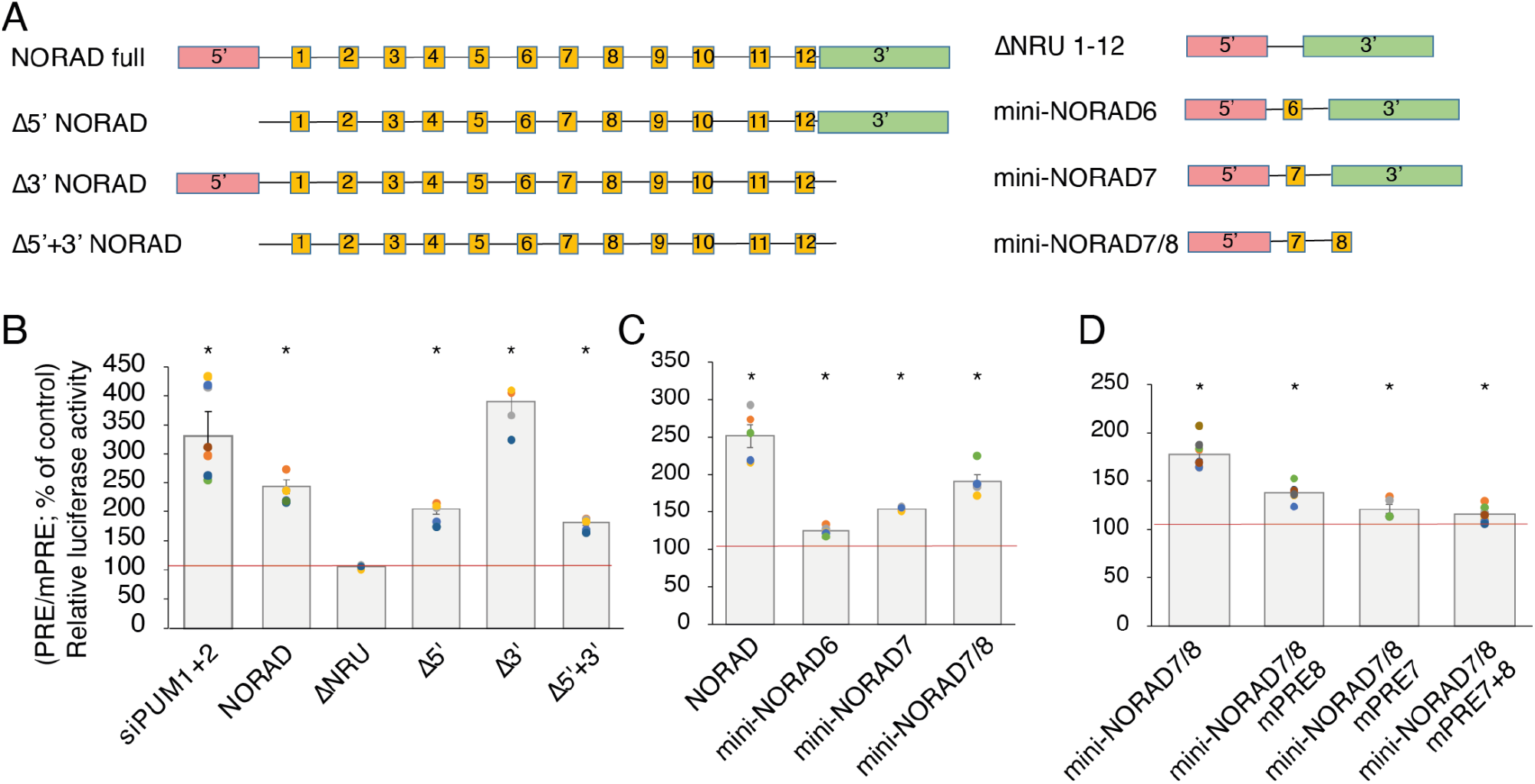
Modular contribution of NORAD sequences to de-repression of a Pumilio sensor. **(A)** Schematic representation of the different NORAD region combinations used for pcDNA3.1-based over-expression in conjunction with luciferase assays based on 8XPRE and 8XmPRE reporters. **(B-D)** Normalized luciferase levels in cells co-transfected with 25nM of specific PUM1, PUM2 siRNAs with PRE-reporter (first column) or over-expressing the sequences presented in A. The data are shown as the percentage change from control, where control is designated 100% (scrambled RNA or empty plasmid - red line). Results are presented as means ± SEM based on at least three independent experiments. Asterisks indicate significant differences from control; (*) P < 0.05, t-test.

We first compared constructs where we removed the 5’ region (bases 1–573, Δ5’ in **Fig. 5A**), a 3’ region (bases 4682–5343, Δ3’ in **Fig. 5A)**, or the middle part of NORAD (bases 604–4774, ΔNRU in **Fig. 5A**) which contains all the PRE clusters. Expression of the WT full-length NORAD resulted in ∼2.5-fold derepression of the 8XPRE reporter compared to 8XmPRE (**Fig. 5B**). As expected, removal of the middle part of NORAD completely abrogated the derepression (**Fig. 5B**). Removal of the 5’ region also had a significant effect, whereas removal of the 3’ region led to a significantly stronger de-repression than that caused by the full-length NORAD (**Fig. 5B**). The combined removal of the 5’ and 3’ modules (Δ5’+3’) led to an effect similar to the removal of the 5’ end (**Fig. 5B**). Notably, some of these changes also affected NORAD expression levels, although to a lesser extent (**Fig. S5B**).

We next tested if a shortened version of the middle part, containing only a subset of the NRUs, is sufficient for de-repression (**Fig. 5B**). Indeed, a combination of the 5’ region with just one of NRU 6 or 7 had a limited effect on 8XPRE levels (**Fig. 5C**), whereas the combination of NRUs 7+8 with the 5’ module (‘mini-NORAD7/8’) was sufficient for potent ∼2-fold de-repression of the reporter (**Fig. 5C**), despite having only 1,443 nt of the 5.3 kb in the full NORAD sequence. As expected, mini-NORAD7/8 activity was abolished when the three PREs in repeats 7 and 8 were mutated (**Fig. 5D**).

### A structured region between the PREs in repeats 7 and 8 contributes to Pumilio repression

We were next interested in the contribution of the extensively paired regions revealed by COMRADES. The region showing the most extensive intra-molecular pairing within NORAD falls between the PRE clusters 5 and 6 (**Fig. 1** and **Fig. 2**), and the predicted fold of the part of NORAD included in mini-NORAD7/8, based on thermodynamics and COMRADES data, suggests that the paired region brings PRE clusters 5 and 6 to be in close spatial proximity to each other (**Fig. 6A**). This paired region is much more conserved in evolution than the ‘loop’ region between the two paired strands, and the predicted structure is supported by low reactivity in DMS-MaPseq data in HEK 293 cell lines (**Fig. 6A-B**, data from (Zubradt et al., 2017)).

**Figure 6.**
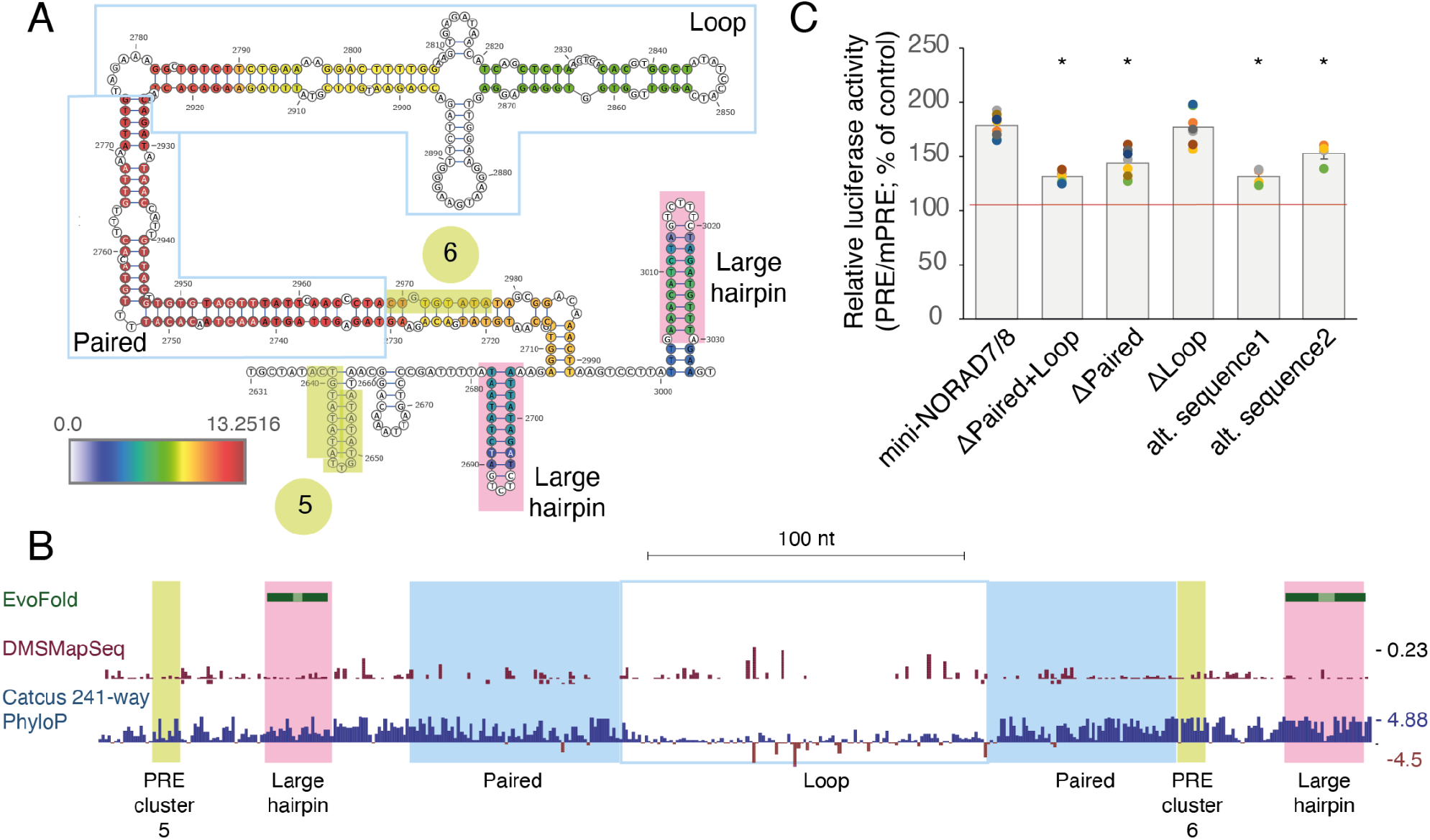
A structured region between PRE clusters 5 and 6 is important for Pumilio derepression. **(A)** Schematic representation of the predicted fold between PRE clusters 5 and 6, color-coded as in **Fig. 3. (B)** Structured regions predicted by EvoFold (Pedersen et al., 2006), DMS-MaPseq reactivity scores (Zubradt et al., 2017) taken from the RASP database (Li et al., 2021) and PhyloP scores (Pollard et al., 2010) computed based on 241-way multiple sequence alignment, taken from the UCSC genome browser hg38 assembly. **(C)** Normalized luciferase levels in cells over-expressing mini-NORAD7/8 variants, where the regions indicated in A are deleted, or where two alternative sequences with the same predicted fold (alt. sequence 1 and 2) were used. The data are shown as the percentage change from control, where control is designated 100% (empty plasmid - red line). Results are presented as means ± SEM based on at least three independent experiments. Asterisks indicate significant differences from overexpression of mini-NORAD7/8 plasmid; (*) P < 0.05, t-test.

We next examined the functional contribution of the structured region in the context of mini-NORAD7/8. Removal of the whole structured region (‘paired+loop’) substantially diminished Pumilio de-repression (**Fig. 6C**), as did independent removal of just the more paired region. In contrast, removal of the loop part of the structure had a limited effect (**Fig. 6C**). We next wondered whether the structure of this region is sufficient for its function. We used RNAinverse (Lorenz et al., 2011) to design two RNA sequences with the same predicted fold as the structured region but a different sequence, and replaced the structure within the context of mini-NORAD7/8. Interestingly, this recoded sequence had a significantly impaired ability to de-repress the reporter (**Fig. 6C**), suggesting that both the structure and the sequence in this region contribute to NORAD function. Interestingly, when the 7/8 hairpin region was deleted in the context of the full NORAD, it did not affect its ability to de-repress the Pumilio reporter (**Fig. S4D-E**), which was likely facilitated by the maintained interactions between the other PRE elements (**Fig. 3**).

### Intermolecular interactions between NORAD and other RNAs

We next focused on RNA-RNA interactions between NORAD and other RNAs. We identified chimeric reads linking NORAD with other RNAs and used DESeq2 (Love et al., 2014) to evaluate the significance of the enrichment in COMRADES samples compared to the controls in which the psoralen crosslinking was reversed before the proximity ligation (**Table S2**). Among non-coding RNAs, only U1 snRNA had a significant and reproducible enrichment of interactions with NORAD, most of which were localized in three regions, and largely unaffected by the Ar treatment (**Fig. S6**). Interactions with mRNAs appeared in other regions, were spread throughout the NORAD locus with a notable peak between NRUs 2 and 3 and in the ‘loop’ region between NRUs 7 and 8, and their pattern was also largely unaffected by Ar treatment (**Fig. S6**).

When grouping the chimeric reads by the interacting protein-coding gene, there were 34, 30, and 12 RNAs with a number of chimeras higher in the crosslinked cells compared to their controls in untreated, Ar-treated, and Doxo-treated cells, respectively (fold-enrichment >1.25 and P<0.05, **Table S2**). Five RNAs were shared between untreated and Ar-treated cells. These transcripts were neither significantly enriched for PREs in their 3’UTRs nor significantly affected by NORAD depletion in HCT116 cells, suggesting that NORAD does not have a preference to basepair in the cell with other Pumilio targets (**Fig. 7A-B**). However, transcripts enriched in each of the conditions were significantly more likely to be enriched in stress granules compared to other genes, with the most significant enrichment observed for transcripts interacting with NORAD in Ar-treated cells (**Fig. 7C**). Notably, there was no substantial difference in the regions of NORAD enriched with chimeric reads with the 2,462 genes enriched in stress granules (Fold-change >2, P<0.05) vs. regions chimeric with segments other genes (**Fig. S5)**, suggesting that there is no particular region in NORAD that preferentially interacts with stress-granule–localized genes.

**Figure 7.**
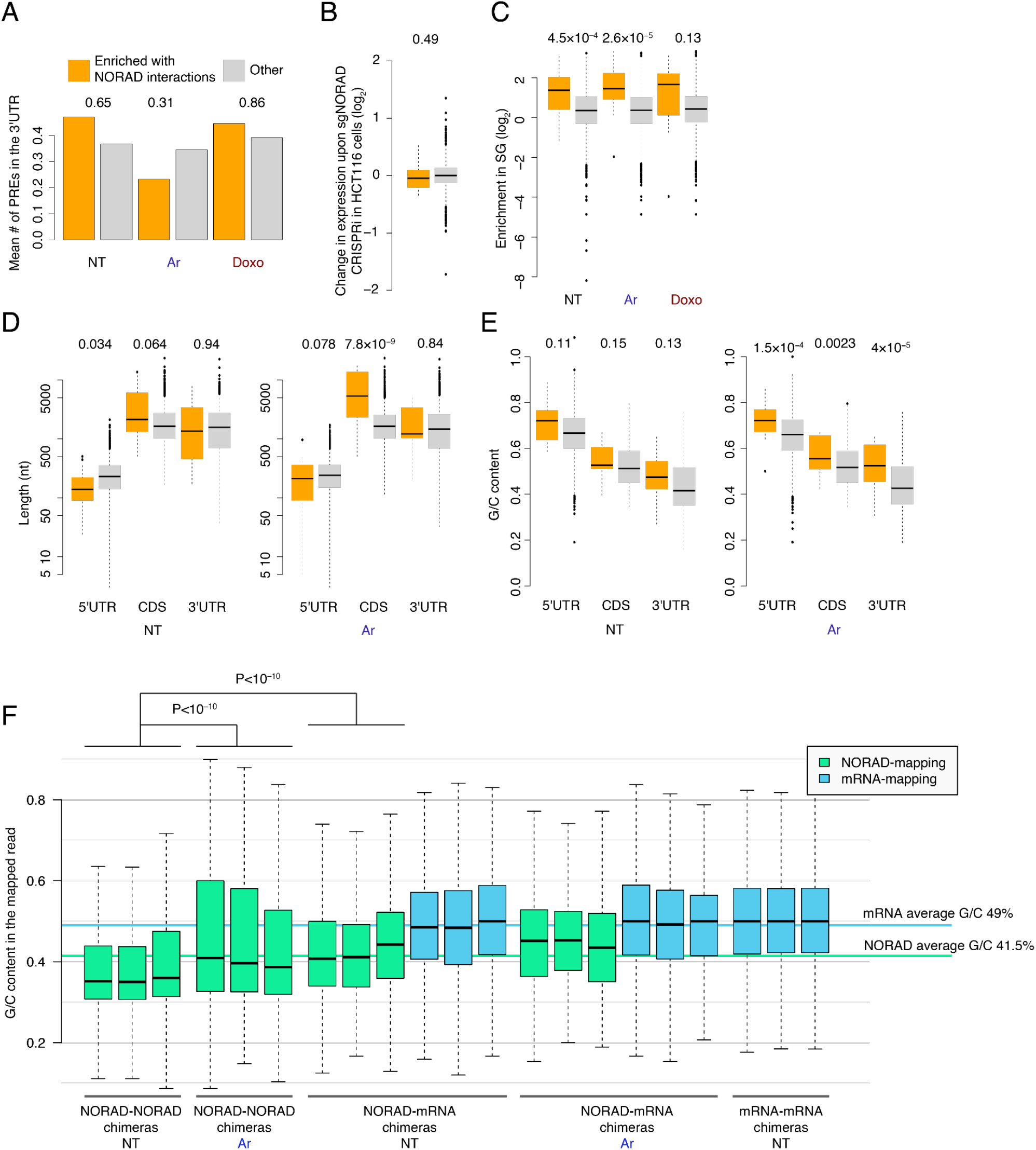
RNA-RNA interactions between NORAD and mRNAs. **(A)** Average number of PREs (defined as UGUANAUA) in the 3’ UTRs of mRNAs enriched in chimeric reads with NORAD in the indicated condition, and all other mRNAs. T-test P-values comparing the enriched and the other genes are shown above each condition. **(B)** Changes in gene expression at 96 hr after CRISPRi-mediated KD of NORAD in HCT116 cells (data from (Munschauer et al., 2018)) for genes enriched in interactions with NORAD in NT conditions and all other genes. Wilcoxon rank-sum test P-value comparing the enriched and the other genes is shown above. **(C)** As in B for enrichment in the stress granule transcriptome, using data from (Khong et al., 2017). **(D)** As in B for the length of the indicated mRNA part, for genes enriched in the NT (left) or Ar (right) condition. **(E)** As in D for the G/C content of the indicated mRNA part. **(F)** G/C content of the part of the read mapped to NORAD or to the mRNAs. Each data point corresponds to a UMI in the sequencing data, and three biological replicates are shown separately in each comparison. Horizontal lines indicate the average G/C content in mRNAs (weighted averages of the 5’UTRs, CDS, and 3’UTRs) and the average G/C content of the NORAD sequence. Comparisons done using Wilcoxon rank-sum test.

As the stress-granule transcriptome was reported to preferentially include genes with specific characteristics, we analyzed the length and the G/C content of the mRNAs enriched in NORAD interactions, and found that they had longer coding sequences (CDS), in particular when considering the enrichments in the Ar conditions, and higher G/C content throughout the RNA (**Fig. 7D-E**). A longer CDS (but not UTRs) has been previously associated with stress-granule enrichment (Khong et al., 2017). We then wondered if the differences in G/C content might reflect a general tendency for COMRADES to recover interactions of G/C-rich RNAs. To test this, we considered all the chimeric NORAD-NORAD and NORAD-mRNA reads, divided them into the fragments that mapped to NORAD and those that mapped to the mRNA, and computed the respective G/C content of each fragment (**Fig. 7F**). The G/C content of the mRNA fragments was similar between NORAD-mRNA hybrids and mRNA-mRNA hybrids in an un-enriched HCT116 COMRADES dataset, and matched the average mRNA G/C content, arguing against a strong bias in COMRADES data. Interestingly, within NORAD, in the NT samples, chimeric reads mapping to different NORAD parts were significantly less G/C rich than those mapping to NORAD and mRNAs, and below the average G/C content of the NORAD sequence. In Ar-treated cells, NORAD-NORAD chimeras were more G/C-rich, fitting the increase in interactions in the G/C-rich 5’ module and the decline in interactions in the more A/U-rich central part (**Fig. 4**).

Intramolecular RNA-RNA interactions of NORAD, enriched in the proximity to the A/U-rich PREs and the A/U-rich SAM68 binding sites we described previously (Tichon et al., 2018), are thus preferentially formed between more A/U-rich regions of NORAD, whereas the intermolecular interactions, presumably mostly occurring at the “outer surface” of the folded NORAD in the cell, are more G/C-rich, less sensitive to Ar stress, and preferentially connected NORAD with long, G/C-rich RNAs that travel to the stress granules upon Ar treatment.

## Discussion

Our structural analysis indicates that NORAD folds hierarchically and modularly, with separate 5’ and 3’ modules and a middle region containing the NRUs each folded into separate modules (structural domains). Furthermore, long-range interactions within the central region help position the PREs in the NRUs in closer spatial proximity to each other than expected by chance, with a particularly strong interaction formed by an extensively paired region between the PREs in NRUs 7 and 8. We used these findings to design a ‘mini-NORAD’ gene that can potently de-repress a Pumilio reporter and is thus instrumental for future studies of the ability of NORAD to repress Pumilio activity.

We suggest that the different modules within NORAD contribute to different aspects of its function. The two 5’ modules are substantially more G/C-rich and more rapidly evolving than the rest of NORAD sequence (**Fig. 4**), and do not contain prominent PREs. The 5’ modules contain the sequences that were previously shown to be sufficient for an NXF1-dependent export of a single-exon RNA from the nucleus (Zuckerman et al., 2020) as well as a region showing extensive interactions with the RBMX protein (Munschauer et al., 2018). This region was also shown to be the least effective in recruiting a reporter RNA to stress granules upon arsenite treatment (Matheny et al., 2020), and we see its structure is least affected by Ar treatment (**Fig. 1**). We find that this region is important for the ability of NORAD to inhibit Pumilio repression, and suggest that it likely does so through ensuring efficient export of NORAD to the cytoplasm and/or its stability there, while potentially being responsible for additional functions in the nucleus through its interaction with RBMX.

The functions of the 3’ part of NORAD remain enigmatic. In our experimental setup, we find that it limits the ability of NORAD to inhibit Pumilio function, potentially by limiting its expression. One potential reason could be that without this region, the PREs in NORAD become physically closer to its poly(A) tail, a feature associated with more efficient repression of Pumilio proteins (Wolfe et al., 2020). It is possible that one function of the 3’ module is thus to provide a “buffer” between the PREs in the NRUs and the poly(A) tail and to limit the ability of the NORAD PREs to induce its degradation. It is also possible that this region, which contains many bases conserved in evolution, also serves additional, Pumilio-unrelated functions.

It was recently shown that NORAD nucleates the formation of phase-separated PUM condensates, termed NORAD-PUM (NP) bodies (Elguindy and Mendell, 2021). Our results complement this model, as they suggest the structure of NORAD helps position some of the PREs in close spatial proximity to each other in a manner that likely increases NP body formation efficiency. For instance, it was demonstrated that four but not two PREs are sufficient for NP formation, but only in a specific context, as mRNA 3’ UTRs with multiple PREs did not efficiently promote NP body formation (Elguindy and Mendell, 2021). mini-NORAD7/8 contains three spatially co-located PREs, and additional U/A-rich sequences that can potentially also bind Pumilio proteins, or proteins that can interact with the Pumilio proteins, such as SAM68 (Tichon et al., 2018). Single NRUs 7 or 8, with just one or two PREs, do not appear to be sufficient for potent Pumilio de-repression that requires both PRE clusters, as well as the paired structured regions connecting them. The structured regions connecting NORAD PREs thus likely mediate its efficient ability to form NP bodies that is absent in Pumilio-bound multi-PRE 3’UTRs. Further, NP-bodies were shown to increase in size upon DNA damage induction, and interestingly, we find that the clustering of the PREs becomes more significant upon Doxo treatment.

We report that Ar treatment, that leads to a prominent shift of NORAD localization to stress granules (Khong et al., 2017; Matheny et al., 2020; Moon et al., 2020) also results in a structural rearrangement within NORAD, which includes a major reduction in intra-molecular contacts within most of NORAD sequence, with a notable exception of the 5’ module, and, to a lesser extent, a reduction in inter-molecular contacts with other RNAs. We cannot exclude the possibility that the stress granule environment is less accessible for the psoralen probing reagent we are using, which may underlie some of these changes. Interestingly, Ar treatment overall did not affect the clustering of the PREs within NORAD, which remained significant (**Fig. 3B**), and correspondingly, NORAD over-expression prior to Ar treatment de-repressed the Pumilio reporter as efficiently as in untreated cells (**Fig. S5F**). The changes in NORAD structure upon Ar treatment thus do appear to impact the ability of NORAD to inhibit Pumilio, but may have other, unrelated, stress-granule–related functions. Notably, we did not observe any large-scale changes in the numbers or morphology of stress granules in NORAD-depleted Ar-treated cells.

One limitation of our study is that whereas we characterized NORAD structure in cells where it is expressed endogenously at physiological levels, for the luciferase reporter assays we use an over-expression setting in WT cells, in which NORAD levels are higher than in the physiological setting. The advantage of this approach is that in U2OS cells it allows us a large dynamic range of up to ∼4-fold de-repression of our sensitive 8XPRE reporter, which is better than what was observed in WT or NORAD^−/–^ HCT116 cells upon NORAD expression ((Tichon et al., 2018) and **Fig. S5G**). Our experimental model thus allows us to effectively measure differences between the different NORAD variants. Notably, several features that we observed in this setting closely match those observed in a stable expression system, such as the requirement of the canonical PREs for NORAD function, and the observation NRUs 7 and 8 are sufficient for suppressing the chromosomal instability resulting from NORAD loss (Elguindy et al., 2019).

A recent study has characterized the folding of the NORAD in vitro by the nextPARS approach, which allows measurements of reactivities of individual bases that can be used to inform structure predictions (Chorostecki et al., 2021). This structure is concordant with our finding that NRUs mostly fold independently, with occasional inter-NRU interactions, in particular between NRUs 1–10. When we originally characterized the 12 NRUs and the similarities between them, we noted that several conserved elements, including a small and a larger hairpin are peculiarly found in some NRUs and not others (Tichon et al., 2016). The nextPARS probing data analysis has suggested that in fact some of the NRUs that do not contain these elements do fold into similar structures, but some of the time (Chorostecki et al., 2021). Specific proteins that recognize and bind these elements have remained elusive so far. Based on our findings here, a sensible hypothesis is that these small structured elements mainly function in the context of the broader ‘task’ of the overall NORAD structure to spatially position the PREs at favorable distances and orientations relative to one another. This can explain the tolerance of NRUs in evolution, where individual NRUs lost some of the elements that were presumably present in the ancestral NRU prior to its duplication.

## Methods

### Cell culture

HCT116 cells were cultured in McCoy’s 5a medium supplemented with 10% fetal bovine serum (FBS) and 100 U of penicillin/0.1 mg mL^−1^ streptomycin. U2OS cells (osteosarcoma; obtained from American Type Culture Collection) were routinely cultured in DMEM containing 10% fetal bovine serum (FBS) and 100 U of penicillin/0.1 mg mL^−1^ streptomycin. All cells were maintained at 37°C in a humidified incubator with 5% CO_2_. and were routinely examined to rule out mycoplasma contamination.

### COMRADES

The COMRADES method was performed as previously described (Ziv et al., 2018). Independent biological replicates were performed using ∼150 million cells each. Ar treated HCT116 cells were supplemented with 20 mM sodium arsenite and maintained for 1 hour under growth conditions. Doxo-treated HCT116 cells were supplemented with 1 uM Doxorubicin and were maintained for 24 hours under growth conditions. Cells were washed 3 times with HANKS buffer and were incubated with 0.7 mg/ml Psoralen-triethylene glycol azide (psoralen-TEG azide, Berry & Associates) diluted in PBS and supplemented with OptiMEM I (Gibco) without phenol red for 20 minutes. Subsequently, cells were irradiated with 50 KJ/m2 365 nm UVA on ice using a CL-1000 crosslinker (UVP). Cells were lysed using RNeasy lysis buffer (QIAGEN) supplemented with DTT and proteins were degraded using proteinase K (NEB). Total cellular RNA was purified using the RNeasy maxi kit (QIAGEN) and quantified using Qubit RNA BR assay kit.

### NORAD enrichment

RNA was mixed with a tiling array of 50 antisense biotinylated DNA probes, 20 nt each (IDT), targeting human NORAD. RNA was maintained at 37C overnight under constant rotation in 500 mM NaCl, 0.7% SDS, 33 mM Tris-Cl pH 7, 0.7 mM EDTA, 10% Formamide. Dynabeads MyOne Streptavidin C1 (Invitrogen) were added, and RNA was maintained for an additional hour rotating at 37°C. Beads were washed 4 times with 2x SSC buffer supplemented with 0.5% SDS and 1 time with 2x SSC buffer without SDS. RNA was released from the beads by degrading the DNA probes using 0.1 units/ml Turbo DNase (Invitrogen) at 37°C for 30 minutes. RNA was cleaned using RNA Clean & Concentrator (Zymo Research) following the manufacturer protocol for capturing RNA bigger than 200 nucleotides.

### Crosslinked RNA enrichment

NORAD-enriched RNA was fragmented by 20 minutes incubation at 37°C with 0.1 units/ml RNase III (Ambion). Reactions were terminated by cleaning RNA with SPRI beads (Amersham) supplemented with Isopropanol. Biotin was attached to cross-linked RNA duplexes by incubating at 37°C for 1.5 h with 150mM Click-IT Biotin DIBO Alkyne (Life technologies) under constant agitation. Following SPRI beads cleanup, biotinylated RNA duplexes were enriched using Dynabeads MyOne Streptavidin C1 (Invitrogen) under the following conditions: 100 mM Tris-Cl pH 7.5, 10 mM EDTA, 1 MNaCl, 0.1% Tween-20, 0.5 unit/ml Superase-In (Invitrogen). Beads were washed 5 times with 100 mM Tris-HCl pH 7.5, 10 mM EDTA, 3.5 M NaCl, 0.1% Tween-20, and RNA was eluted by adding 95% Formamide, 10 mM EDTA solution preheated and incubating at 65°C for 5 minutes. RNA was purified using RNA Clean & Concentrator (Zymo Research). Each RNA sample was split in two: one half was proximity ligated, following UVC irradiation to reverse the crosslink (i.e., interactions sample). The other half was UVC irradiated to reverse the crosslink, and only then was proximity ligated (i.e., control sample). Prior to proximity ligation, RNA was denatured by heating to 90°C and transferring to ice water. Proximity ligation was performed using 1 unit/ml RNA ligase 1 (NEB), 1x RNA ligase buffer, 50mM ATP, 1 unit/ml Superase-in (Invitrogen), at a final volume of 200ml. Reactions were incubated overnight at 16°C and were terminated using RNA Clean & Concentrator (Zymo Research). Crosslink reversal was done by irradiating the RNA with 2.5 KJ/m2 254 nm UVC using a CL-1000 crosslinker (UVP) on ice. Sequencing library preparation was done as described in (Ziv et al., 2020). Paired-end 100nt libraries were sequenced using HiSeq 1500 sequencer (Illumina).

### COMRADES data analysis

After obtaining sequencing data in FASTQ format, we removed sequencing adapters (cutadapt -a AGATCGGAAGAGCACACGTCTGAACTCCAGTC -A AGATCGGAAGAGCGTCGTGTAGGGAAAGAGTGT -- minimum-length 10), merged paired-end reads using pear with default settings, collapsed identical reads (bash uniq -c), extracted the 6-nt Unique Molecular Identifier (UMI) from 3’ends of reads, and saved the reads in a deduplicated fasta file, where the number of duplicates and number of UMIs observed for every sequence was encoded in the read ID. To call chimeric reads, we used hyb (Travis et al., 2014) with bowtie2 mapping to a human transcriptome database (Helwak et al., 2013) that consisted of spliced mRNAs, tRNAs and other noncoding RNAs, supplemented with the human NORAD sequence (nucleotides 1-5339 from transcript NR_027451.1). We also performed the mapping against a database that consisted of the NORAD sequence alone, which recovered approximately 10% more NORAD:NORAD chimeras, without noticeably altering the pattern of interactions.

We obtained 2M-11M unique mapped reads per replicate experiment, of which 3–4% were chimeric reads; in the untreated and Doxo experiments, 0.8–1.3% of all mapped reads were NORAD:NORAD chimeras, while in Ar experiments, 0.2–0.3% of reads were NORAD:NORAD chimeras. In the control experiments, in which the order of UVC irradiation and proximity ligation was reversed, the fraction of NORAD:NORAD chimeras ranged from 0.02% to 0.12%. Between 3 and 19% of all mapped reads, chimeric or non-chimeric, were mapped to NORAD in the NT, Ar, and Doxo datasets. By contrast, 0.002% of reads were mapped to NORAD in a control COMRADES experiment that did not include NORAD pulldown.

Pearson correlations between replicate experiments were calculated using numbers of NORAD:NORAD chimeras in 100 nt x 100 nt windows, and were plotted in R using the “corrplot” package. The same data was used for the hierarchical clustering of the experiments. 2-dimensional maps of chimera coverage were generated using Java Treeview. To calculate topological domains alongside NORAD, we used the coverage of chimeras in 10 nt x 10 nt windows as an input matrix for TopDom (Shin et al. 2016), and ran TopDom with window.size=30, which resulted in an effective window size of 300 nt. The other TopDom settings were kept as default.

Regions with significantly distinct NORAD:NORAD interactions between the 3 NT datasets and 3 Ar datasets were calculated using DESeq2 based on the numbers of chimeric reads in 10×10 windows (Love et al., 2014). The direction of change was distinguished using log fold change values, and –log_10_ p-values (capped at 2/–2) were used to plot the interaction heatmap in R, where color intensity reflects statistical significance. Example interactions were isolated and plotted in R with normalization of chimeric counts per million mapped reads, alongside their predicted structures.

Arc plots (**Fig. 3**) were plotted in R with the “R4RNA” package based on the R-chie web server (Lai et al., 2012). The color of arcs represents the p-values, which were calculated by counting the interactions between two connected PRE sites with a 50nt radius (e.g. between PRE1:710–817 and PRE2:1087– 1195), and comparing it to 50nt circular permutations along the genome (e.g. 760–867 and 1137–1195, and so on) using independent t-tests. The resulting p-values were then subjected to Bonferroni correction before plotting.

NORAD secondary structures were predicted using the comradesFold pipeline (Ziv et al. 2018), and the structures were plotted in VARNA. comradesFold generates a set of folding constraints from hyb files, which is then shuffled randomly 1,000 times, and used for structure prediction using hybrid-ss-min. The whole NORAD secondary structure was generated as 6 parts: 1–760, 761–2050, 2051–2630, 2631–3400, 3401–4150, and 4151–5339, each combining domains determined by the TopDom algorithm, with slight adjustments to preserve high confidence structures. Color-coding of base-pairing is based on hyb data-based supporting reads, and was normalized to log2 chimeric counts per million mapped reads.

Numbers of interactions between PRE clusters to other RNA fragments, and non-PRE fragments to other RNA fragments (Figure S3A) were calculated using 50 nt radius for PRE clusters, and 116 randomly generated 110nt-long non-PRE fragments. The average interaction counts for the 116 non-PRE to other RNA fragments were used to provide sufficient data. The counts were also normalized as chimeric reads per million mapped reads.

COMRADES data has been deposited in the GEO database under the GSE188445 accession (reviewer token kdulcoqehngdfkv).

### Plasmids and siRNAs

Plasmid transfections were performed using GenJet In Vitro DNA Transfection Reagent (SignaGen Laboratories). To overexpress NORAD, the pcDNA3.1 vector previously described (Tichon et al. 2016) and available on AddGene (AddGene #120383 was used). Derivatives and mutations of *NORAD* were prepared by Restriction Free cloning (Unger et al., 2010). Primer sequences are given in **Table S3**. As controls for the overexpression experiments, we used empty pcDNA3.1 (+) vector (Invitrogen). In transfections, 200 ng was used per 30,000 cells in 24-well plates for 48 h before cells were harvested. For luciferase experiments, we generated vectors with 8 WT PREs and mutated PRE reporters as controls. To construct these reporters, eight wild-types PRE repeats (GAAAAT<u>TGTATATA</u>AATCAA) or eight mutated PREs (GAAAAT<u>ACAATATA</u>AATCAA) were inserted using XhoI and NotI sites in 3’ UTR of Renilla gene in the psiCheck-2 dual luciferase reporter vector (Promega). The reporter in the amount of 25 ng and plasmids with different NORAD variants in the amount of 200 ng were introduced per 30,000 cells in 24-well plates. For co-transfection of siRNAs together with reporters, we used LipoJet In Vitro DNA and siRNA Transfection Kit (SignaGen Laboratories), according to the manufacturer’s protocol. Cells were transfected with 25 nM SMARTpool siRNA (Dharmacon) targeting PUM1 and PUM2 sequences (**Table S4**) or scrambled siRNA with 25 ng of reporter for 48 hr before harvesting. For the Ar treatment experiment, U2OS cells expressing reporter and NORAD plasmids were treated with 0.5 mM sodium arsenite for an hour before harvesting.

### Real-time PCR analysis of gene expression

Total RNA was isolated using TRI reagent (MRC), followed by reverse transcription using an equal mix of oligo dT and random primers (Quanta), according to the manufacturer’s instructions. Real-time PCRs were performed using the AB quantitative real-time PCR system ViiA 7 (Applied Biosystems). Fast SYBR Green master mix (Life, 4385614) was used for qPCR with gene-specific primers (**Table S5**). All genes expression levels are presented relative to their relevant control (ΔCt) and normalized to *GAPDH* expression levels (ΔΔCt).

### Luciferase assays

Reporter gene activity was measured as previously described (Van Etten et al., 2013). Briefly, 30,000 cells were plated in a 24-well plate. After 24 hr, cells were co-transfected with psiCheck2 plasmids and indicated NORAD plasmids (as described above). Luciferase activity was recorded 48 h post-transfection using the Dual-Glo Luciferase Assay System (Promega) in the Microplate Luminometer (Veritas). A relative response ratio, from RnLuc signal/ FFLuc signal, was calculated for each sample. The percent of change presented is relative to the control plasmid.

### Statistical analyses

Data are presented as an average ± SEM; experiments were repeated at least three times. Statistical analyses were performed using Student t-test or one-way ANOVA, followed by Bonferroni multiple comparison tests, when appropriate. In all analyses, a value of P < 0.05 was considered significant. Asterisks represent significant differences from their respective controls.

## Data availability

**Table S1**. Pairs of PRE clusters are connected by RNA-RNA interactions located in close proximity to both motifs.

**Table S2**. Interactions of NORAD with mRNAs.

**Table S3**. Primer sequences for Restriction Free cloning of NORAD.

**Table S4**. siRNA target sequences.

**Table S5**. Primer sequences for Real time PCR.

## Supplemental Figures

**Figure S1.**
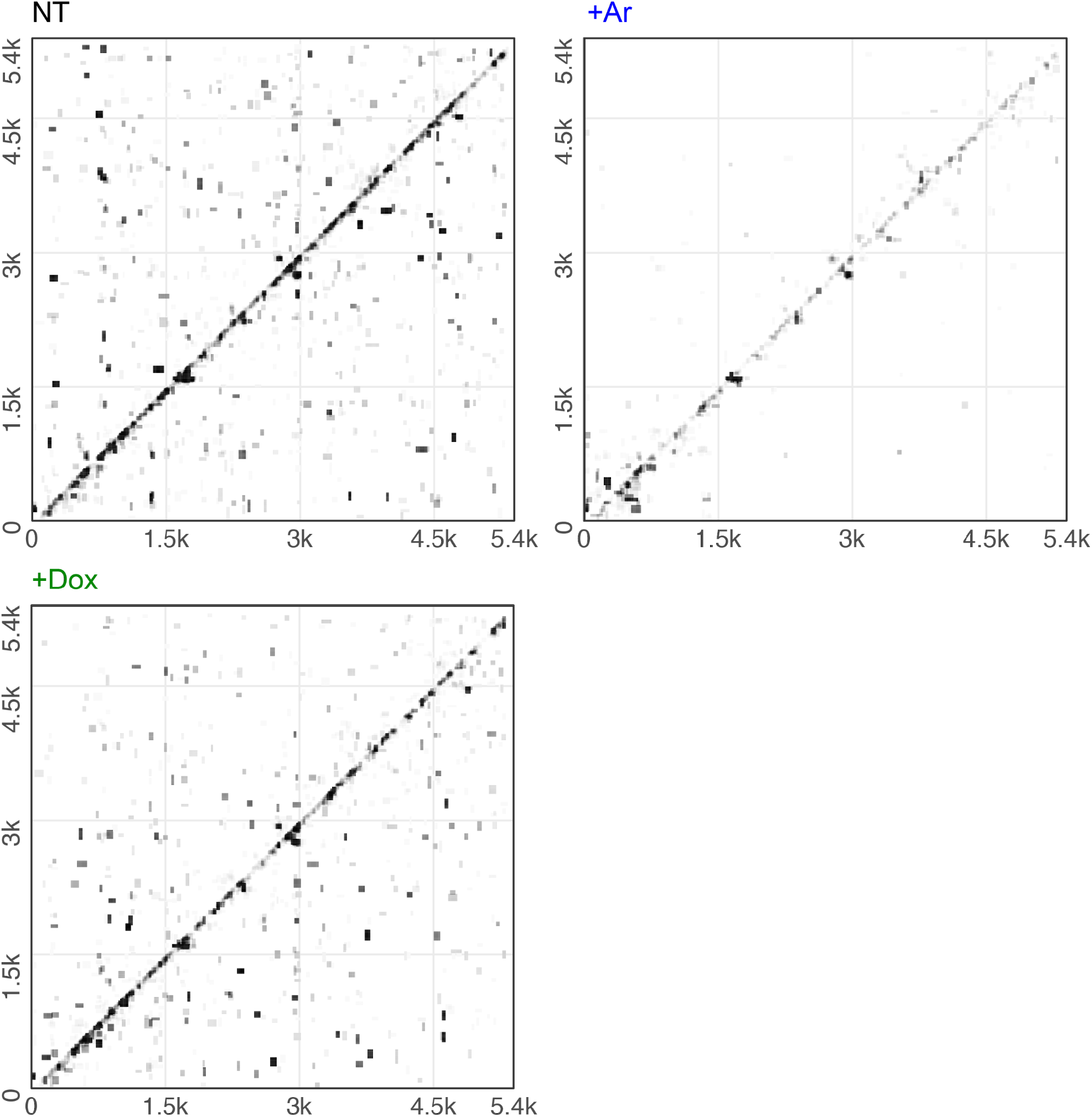
Intramolecular interactions in NORAD in a control experiment. As in Figure 1B, for a control experiment in which the order of UVC irradiation and proximity ligation was reversed.

**Figure S2.**
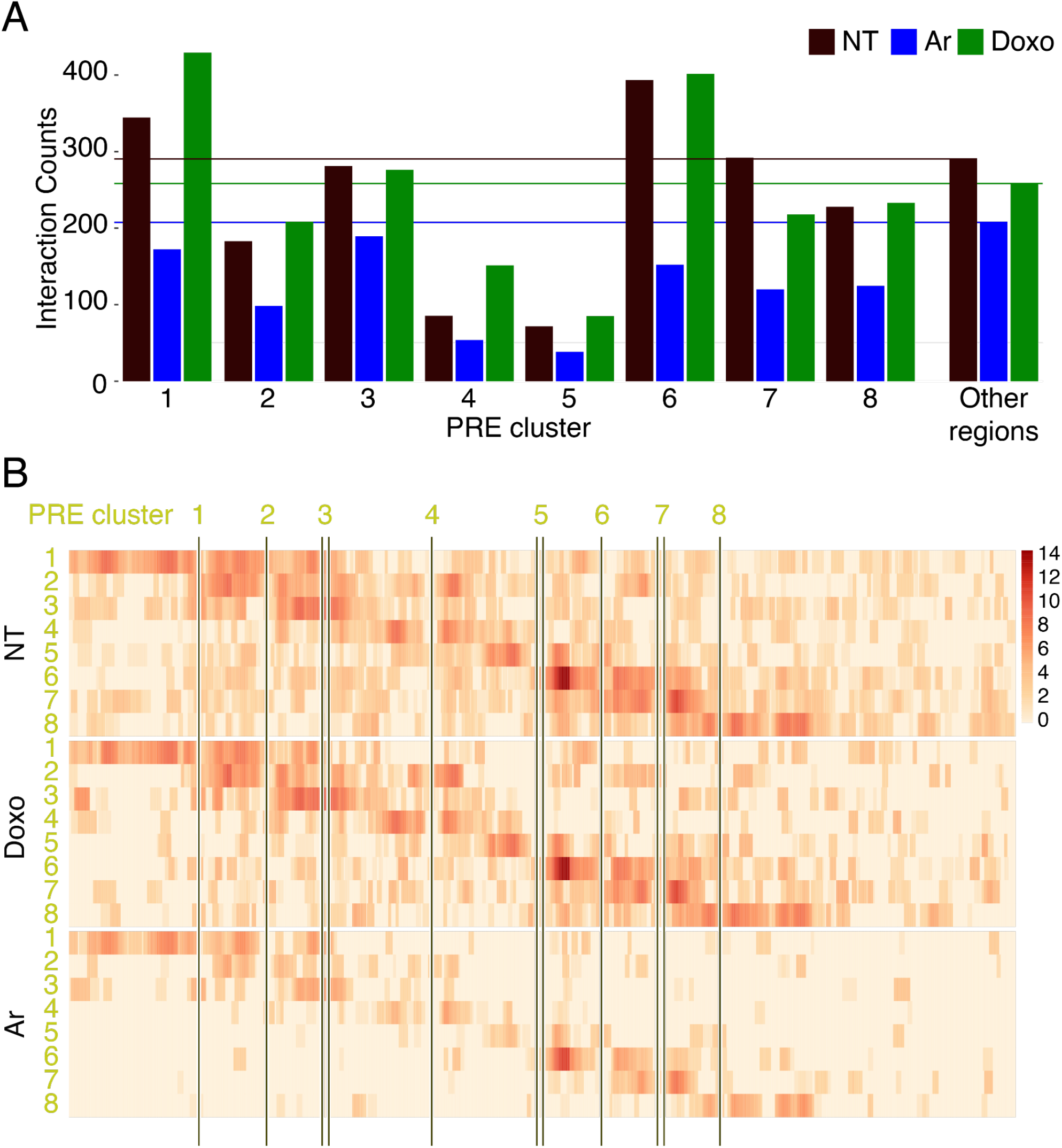
Analysis focused on PRE clusters. **A**. Normalized interaction counts of each of the 8 PRE clusters with a 50 nt radius to other RNA fragments, and other non-PRE regions of NORAD to other RNA fragments, separately for each treatment. Interaction counts for other regions were calculated using the mean of 116 random 110 nt-long non-PRE fragments. Horizontal lines indicate other regions’ mean interaction counts for each treatment. **(B)**. log_2_-transformed normalized numbers of chimeras formed by a region of 50 nt radius surrounding each of the eight PRE clusters and each other based in NORAD. For each indicated treatment, all the biological replicates were combined.

**Figure S3.**
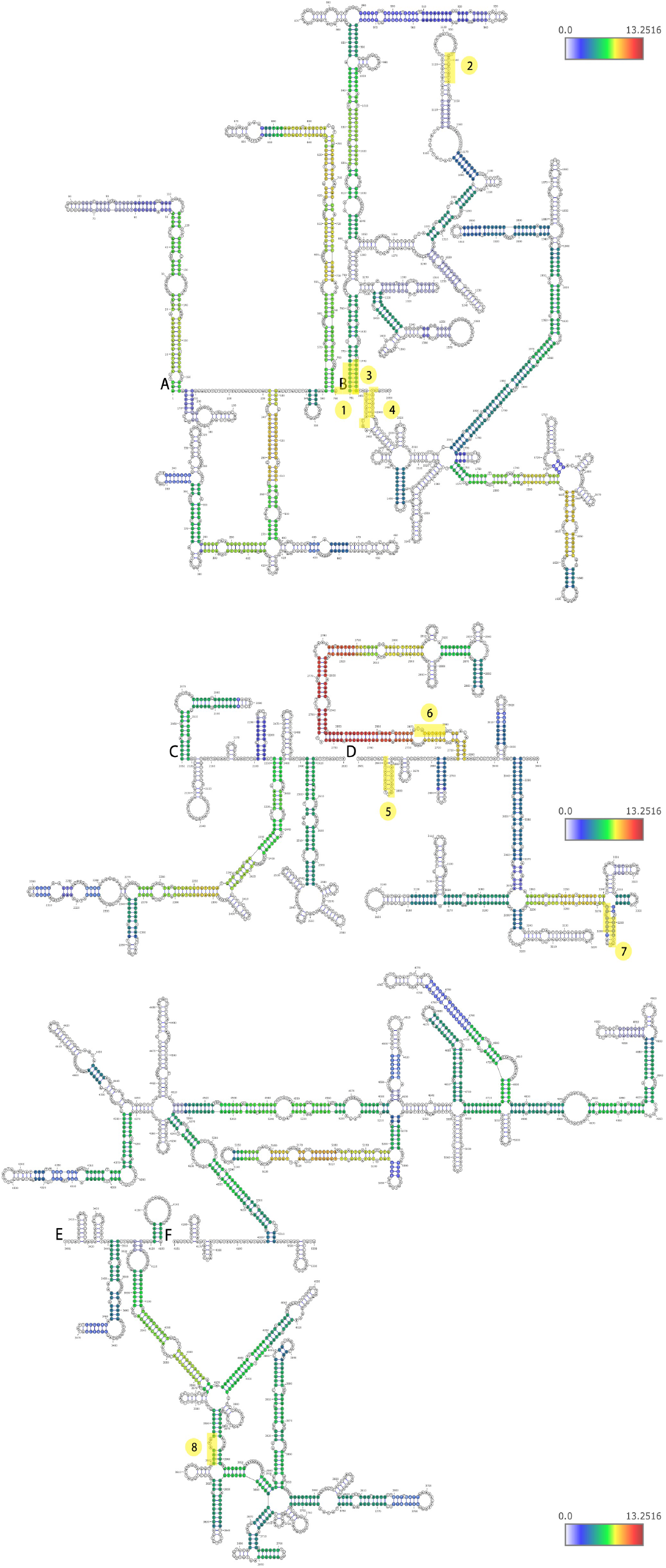
Predicted structures of NORAD in Ar-treated cells. As in Figure 3, for Ar treatment.

**Figure S4.**
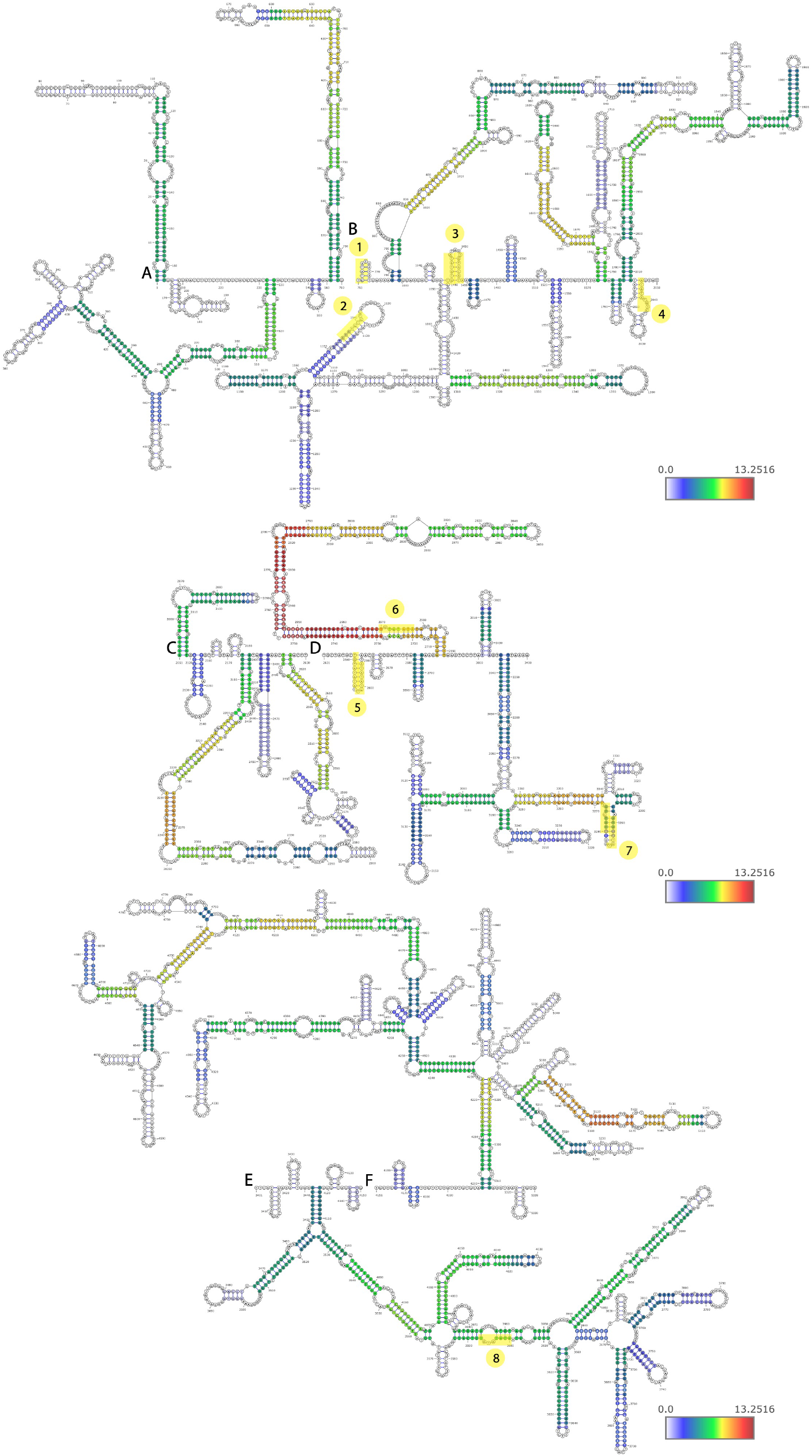
Predicted structures of NORAD in Doxo-treated cells. As in Figure 3, for the Doxo treatment.

**Figure S5.**
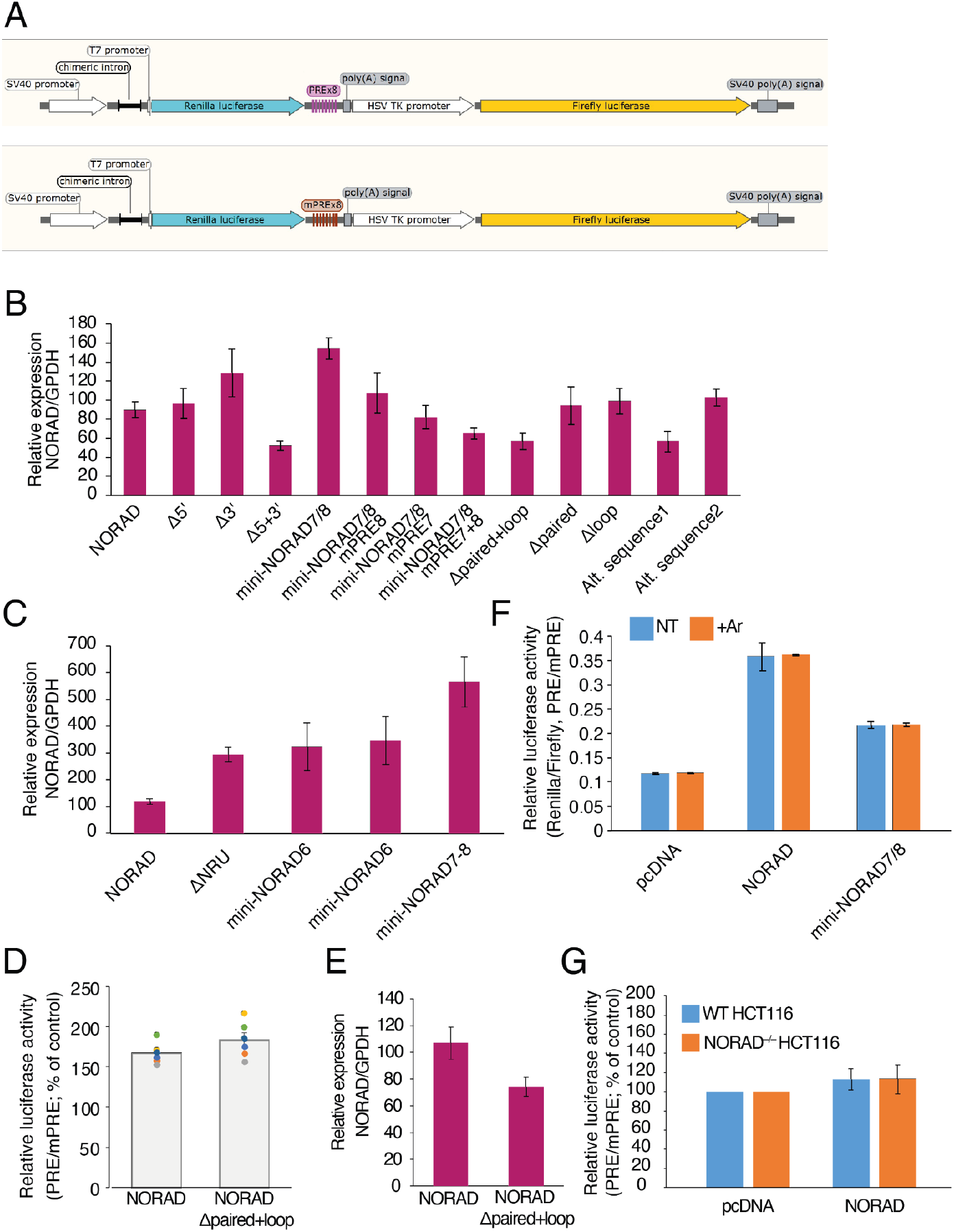
**(A)** Schematic view of the luciferase reporter plasmids used in this study. (**B-C)**. Expression of NORAD as measured by qRT-PCR. Data are presented relative to expression of empty plasmid and normalized to GAPDH expression levels. **(D)** Normalized luciferase levels in cells over-expressing the full length NORAD or full NORAD with a deletion of the paired and loop regions in the NRU7/8 region. The results are presented as means +/- SEM based on at least three independent experiments. **(E)** Same as D for NORAD expression. **(F)** Luciferase reporter results after transfection of the indicated plasmid in cells followed by no-treatment or an Ar-treatment. **(G)** Luciferase reporter results in WT or NORAD^−/–^ HCT116 cells following transfection of the indicated plasmid.

**Figure S6.**
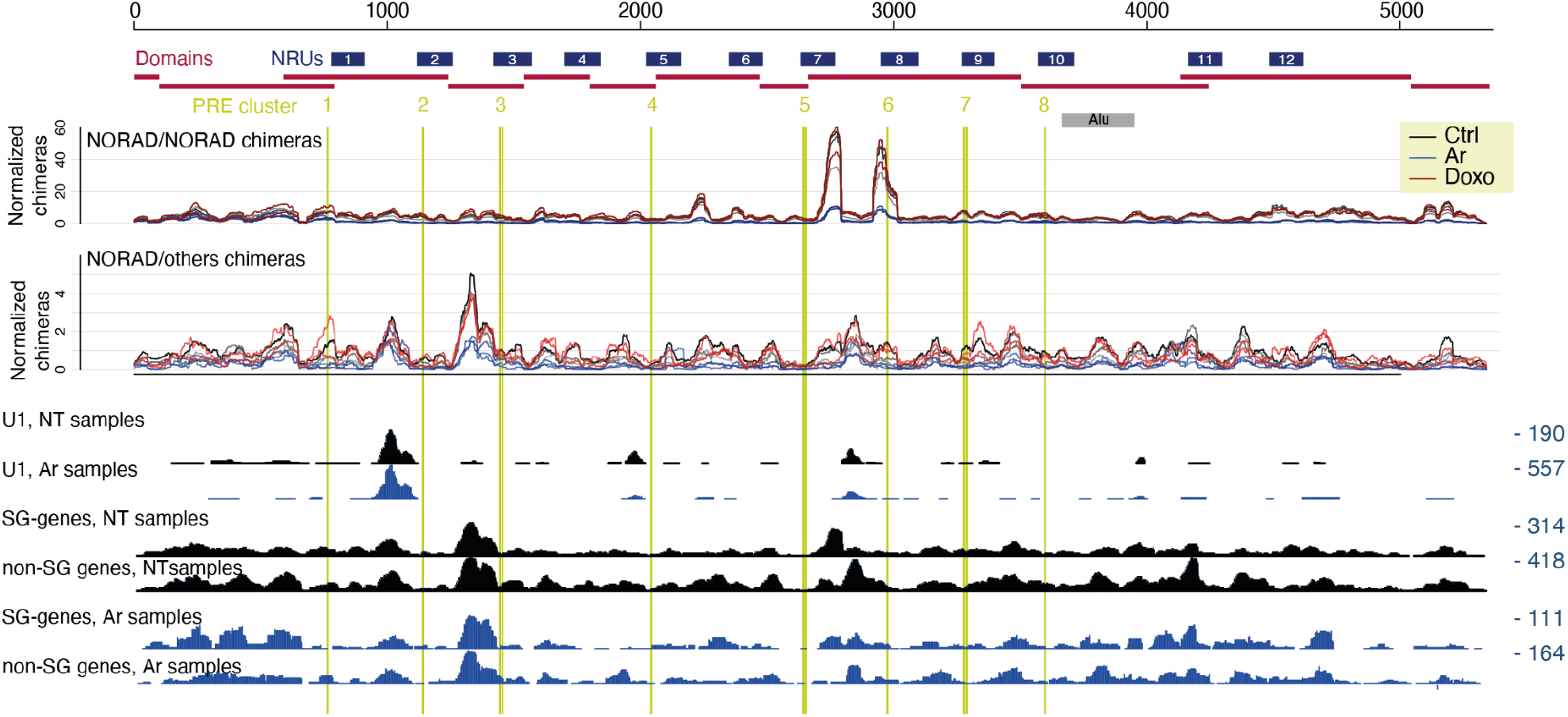
Coverage of chimeric reads with other RNAs across NORAD. The top part is as in Figure 4. At the bottom shown is the coverage of reads chimeric with parts of U1 in NT and Ar-treated samples, coverage of reads chimeric with genes enriched in stress granules (2-fold, P<0.05 enrichment) from (Khong et al., 2017) and with all other genes, in NT and Ar-treated samples.

